# Sequence-Only Prediction of Antibody Fab Thermostability Using Protein Language Model Embeddings

**DOI:** 10.64898/2025.12.30.697053

**Authors:** Michael A. Sennett

## Abstract

Antibody developability is a critical consideration in therapeutic manufacturing, with conformational stability—measured by melting temperature (T_m_)—being a key determinant of manufacturability. Traditional experimental approaches to assess T_m_ are costly and low-throughput, while statistical models offer limited predictive power and generalizability. In this study, we present a scalable, sequence-only machine learning (ML) approach to predict the thermostability of antigen-binding fragments (Fabs) using embeddings from IgBERT, a protein language model fine-tuned on antibody sequences. We trained a Random Forest regressor on 133 proprietary antibody sequences with measured Fab T_m_ values, achieving a Pearson correlation coefficient (PCC) of 0.77 on a held-out internal test set. When applied to a Jain set of clinical-stage therapeutics, the model retained predictive power (PCC = 0.28), outperforming baseline statistical and sequence-similarity methods. Embedding proximity in t-SNE space correlated with prediction accuracy, suggesting that model generalizability is influenced by distributional similarity rather than sequence identity. Our results demonstrate that PLM-derived embeddings encode biophysical features relevant to Fab thermostability and offer a fast, interpretable, and scalable route to de-risk antibody lead candidates.

## Introduction

Predicting antibody developability properties is of interest because it helps to de-risk the manufacturing process. Developability properties, include titer, colloidal stability, conformational stability, poly-specificity, and post-translational modifications.^1,2^ Most of these properties can be measured experimentally for a set of lead candidates. The challenge is that experimentally measuring these properties is not fast, cheap or scalable. Therefore, we cannot apply this approach to a large panel of lead candidates. This challenge has been addressed by developing *in-silico* models to calculate attributes of lead candidates that correlate to poor developability properties or, in limited cases, directly predict developability properties.^3–10^

While all developability properties are important to understand, one particular biophysical property of interest is the melting temperature. Melting temperature is a measure of conformational stability of an antibody. A conformationally stable molecule is necessary because a typical manufacturing process occurs over the course of weeks at ambient temperatures. Therefore, to produce a viable drug in meaningful quantities the antibody must be able to remain folded over the course of the manufacturing process.

Multiple approaches have been used to predict the difference in thermostability across a diverse set of antibody sequences with limited success.^9,11–14^ The most successful approaches to date include a mixture of rational physics-based and machine learning (ML) models. For example, AbMELT is able to differentiate antibodies based on thermostability using a ML model that is trained on molecular dynamic (MD) features.^9^ However, given the computational expense of this approach, it is not scalable when dozens or hundreds of lead-candidates need to be differentiated based on conformational stability.

The question remains; how do we quickly and cheaply identify a set of antibodies most likely to remain folded during the manufacturing process? One answer is to use a model to predict antibody thermostability solely from sequence. Statistical models (e.g., covariance analysis, direct-coupling analysis, and mutual information) are indirect methods that are used to calculate a feature that tends to correlate to antibody thermostability. These models identify conserved residue pairs have been shown to correlate with protein fold contacts, function, and stability with limited success.^15–20^

Despite different implementations, fundamentally these methods rely on the same hypothesis to differentiate proteins by stability; if two residues at two different sites in a protein are in direct contact then introducing a chemically different variant at one of those sites is likely to disrupt the residue-residue contact. We infer that disrupting residue-residue contacts will decrease conformational stability. Though the model is interpretable and fast there is one limitation. There is no scenario in which the introduction of a variant, not normally observed at a pair of covarying sites, would be beneficial to improving the conformational stability of a protein. Hence, we need a model that can directly predict antibody thermostability rather than a feature that may correlate to thermostability under a limited set of conditions.

A ML model would address the two limitations of statistical models. First, a ML model can directly predict thermostability, rather than an attribute correlated to thermostability. Second, there is no inherent constraint that the introduction of a variant will always be detrimental to thermostability. One drawback of implementing an ML model to predict thermostability is that it requires many antibody sequence data labelled with a corresponding melting temperature. However, this can potentially be mitigated by utilizing a transfer learning approach with a protein language model (PLM).^21,22^

PLMs are expected to learn contextual information beyond pairwise interactions and output embeddings that contain features relevant to the biophysical and biochemical nature of a protein.^22–30^ Furthermore, PLMs have been fine-tuned specifically on antibody variable fragments (Fvs) to enhance sequence recovery of Fvs.^31^ The embeddings output from an Fv fine-tuned PLM is expected to encode information specific to Fv structure and function.^6,9,11,13,32–34^ Implementing a PLM reduces the burden of a downstream ML model to learn contextual information about the protein.

A successful model in this case is one that has the ability to differentiate antigen binding fragment (Fab) thermostability for a diverse set. For a model to succeed in this task the prediction error for many sequence mutations need only be accurate over the course of many predictions. This is because the model may incorrectly predict the magnitude and effect of a single variant of a Fab but, on average, the error in predictions for several variants together cancel. In other words, we may accept a model is imprecise in the prediction of the thermostability of a single Fab sequence if it is able to distinguish between thermostable and unstable from a diverse set of Fabs. What is and is not thermostable is somewhat a matter of subjectivity. However, we define, in this case, thermostable Fab as one that has a melting temperature above the average melting temperature of an IgG1 crystallizable fragment (Fc).

In this paper, we establish a Fv sequence-only approach to differentiate the Fab of antibodies on the basis of thermostability. To do this we train a linear regression model utilizing IgBERT embeddings of a diverse internal dataset of antibody melting temperatures. Our model predictions for a held-out subset of our internal dataset have a Pearson correlation coefficient of 0.77 with the ground truth. Although the correlation dropped to 0.28 when extended to a large and dissimilar Jain set of clinical stage therapeutics, our ML model still outperformed our benchmark statistical model approach. By constraining our predictions based on Euclidean distance in t-SNE embedding space we increase the Pearson correlation coefficient to 0.48. This demonstrates the feasibility of training a model to predict Fab melting temperatures based solely on PLM embeddings. This methodology provides a fast and scalable route to differentiate lead-candidates based on conformational thermostability.

## Method

### Data set information

There are a total of 3 different data sets used in this study. Two are proprietary, while the third is from a public source. Our first, and smallest data set, contains 31 sequences from a phage-display panning experiment of our proprietary J.HAL Fabs against SARS-CoV-2 receptor binding domain.^35,36^ The second, internal proprietary data set, used to train our ML model consists of 140 sequences. This set will be referred to as the internal set going forward. Seven of the 140 sequences were held-out for validation. The remaining 133 sequences are subsequently labelled as the training set. Finally, the third data set, contains 133 sequences from a published study by Jain et al.^1^ We will refer to the published set as the Jain set or test set from here on out.

### Identifying Fab T_m_

Our proprietary data sets contain the melting temperature (T_m_) for both the Fc and Fab domain obtained by differential scanning fluorimetry (DSF) in PBS. If there were two T_m_s and if the value of the first T_m_ was below 69 °C, then we used the T_m_ below 69 °C as the Fab T_m_. If there were two T_m_s and if the value of the second T_m_ was above 71 °C, then the second T_m_ was above 71 °C was used as the Fab T_m_. If there was one T_m_, then that was considered to be the Fab T_m_.

### Identifying covariance violations

As outlined by Gunasekaran et al, covariance analysis is a statistical based method to evaluate all possible residue pairs in a given reference multiple sequence alignment.^15^ We implement a similar method with the exception that amino acids-along with a gap character-are divided into 6 different classifications based on the chemical property of the amino acid side-chain rather than two, hydrophobic and hydrophilic. These 6 groups are acidic, basic, hydrophobic, aromatic, neutral polar, and deletions. For a given sequence, a site is considered a violation if chemical property of that residue differs from the expected chemical property based on the same site in the reference alignment. For example, if two covarying hydrophobic sites are normally present in a reference alignment and the sequence of interest is observed to have an acidic residue then it would be considered a covariance violation. The total number of covariance violations is summed over both the heavy and light variable domains.

### Generating IgBERT embeddings to predict biophysical properties

IgBERT is a PLM fine-tuned on unpaired and further fine-tuned on paired antibody sequences from OAS.^31^ For a given pair of sequences IgBERT outputs a set of embeddings. A global average of the output embeddings over the length of the input protein was generated. Consequently, each Fv sequence has the same number of embeddings, 1024. To perform a Zero-shot regression analysis of the output embeddings and the melting temperature, the embeddings were again averaged to reduce the features from 1024 to 1.

In order to train a regressor, the 1024 were reduced in two steps. First, for each embedding a Pearson correlation coefficient with the melting temperature was calculated. Only those embeddings with a correlation coefficient > |0.5| were kept as possible input features. Second, a Pearson correlation coefficient among the remaining embeddings was calculated. Those that had a value < |0.95| were kept as possible features to train a regressor model.

### Regressor training and test

A Random Forest regressor from scikit-learn was trained on 177 embeddings output from IgBERT from 133 of the 140 antibodies in our internal dataset. The resultant model predictions and ground truth were used to calculate the Pearson correlation coefficient, a slope, and intercept. The training and validation was repeated with the SGD and SVM regressors. The Random Forest regression model performed best based on Pearson correlation coefficient, a slope, and intercept of the remaining 7 sequences of the held-out test. Finally, training and validation were again repeated with different hyperparameters for the Random Forest Regressor. The number of trees, the function to measure the quality of a split, and the max features were all varied.

Three models trained with our internal dataset were selected to predict the melting temperature for Jain set. The first model using default hyperparameters in scikit-learn, which used 100 trees, the squared-error cost function, and 100% of the 177 embedding features. The second used 10 trees, the absolute error cost function, and 50% of the embedding features. The third model used 10 trees, the mean squared error cost function, and 50% of the embedding features. For each set of predictions, a least-squares linear regression was performed to calculate the Pearson correlation coefficients, the slopes, and the intercepts.

### Feature reduction

The input features used to train our models were reduced using t-SNE implemented in scikit-learn, using default parameters.^37^ The analysis was performed using the filtered set of IgBERT embeddings from both the internal set and the Jain set.

### Model Metrics

A linear least-squares regression is used to evaluate the true protein sequence T_m_ as a function of model output (e.g, number of covariance violators, sequence log-probability, or predicted melting temperature). For each set of predictions, the Pearson correlation coefficient, the slope, and the intercept were used to assess model performance.

## Results

### Our benchmark statistics-based model correlates to the Fab T_m_ of our internal datasets

Dozens or more potential therapeutics can be isolated from a single discovery campaign against an antigen target. Therefore, computationally scalable approaches are required to differentiate antibody lead candidates based on thermostability. One approach we employ to accomplish this task is a statistical analysis that identifies covarying sites in a given set of sequences.^15^ If the chemical property of a residue is present at one of a pair of sites that is not typically observed then it is flagged as a violation. In limited cases, we observe a correlation between T_m_ and number of covariance violations (Fig. 1, Table 1).

**Table 1:**
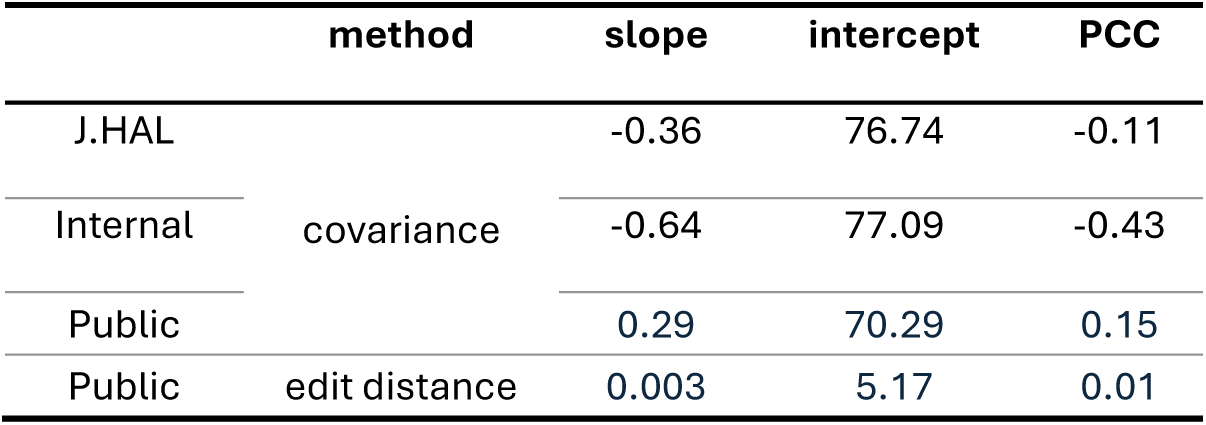
Comparison of molecule property prediction using the number of covariance violators or edit distance for three sets of antibodies.

**Figure 1:**
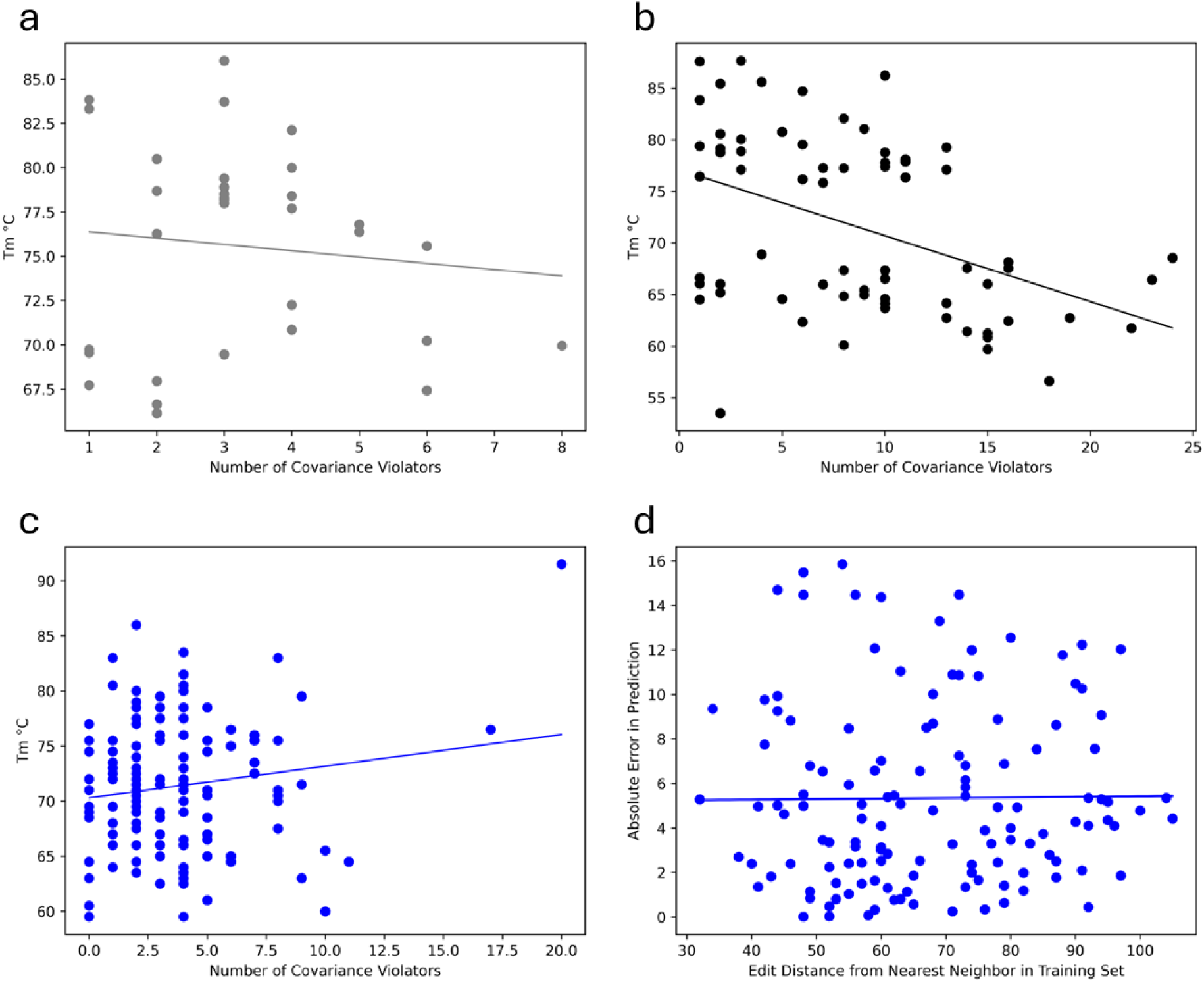
Covariance analysis displays a limited correlation to thermal stability, (a) Plot of antibody melting temperature as a function of the number of covariance violations for Fabs from our AI-generated antibody library, (b) The same as (a), but of biologically derived molecules from our internal dataset, (c) The same as (a), but for a publicly available dataset, (d) Plot of absolute error in melting temperature prediction as a function of the distance of a publicly available sequence from the nearest neighboring sequence in our internal dataset.

Our previous discovery campaign contains a set of molecules generated from an ML model trained to output human-like antibody Fv sequences.^35^ Additionally, we have an internal dataset consisting of antibodies from biologically derived sources with associated T_m_ measurements. The slope of the linear least-squares regression analysis between T_m_ and number of covariance violators is negative, which matches our expectation that disrupting residue couplings should destabilize the protein (Fig. 1a & 1b). Our expectation holds for naturally-derived and artificially-derived Fabs.

How generalizable is covariance analysis? We extended our analysis to the Jain set and plot the T_m_ as a function of number of covariance violations (Fig. 1c). There is a weak correlation among the data, which is demonstrated by Pearson correlation coefficient of 0.15 (Table 1), but crucially, we see the opposite of the expected trend with respect to slope. There is a slight positive slope: the implication is that increasing the number of violations should improve stability when we expect the opposite. This observation demonstrates a limitation of using a covariance analysis.

### A sequence only approach seems to not be correlative to melting temperature

Our statistical model has limited generalizability to melting temperature predictions. However, a reasonable hypothesis would be to assume the melting temperature for one protein is approximately the melting temperature of its closest known homolog. To test this hypothesis, we plotted the absolute difference in melting temperature for the publicly available sequences as a function of edit distance (the number of mutations required to change one sequence to another) from a homolog with a known melting temperature in our internal dataset (Fig. 1d). There is no observed correlation. Predicting melting temperatures using sequence identity only from our internal set is not enough to be predictive of antibody melting temperature.

### An ML model is trained to predict Fab T_m_ only utilizing PLM embeddings

A statistics-based model or sequence-only based model appears insufficiently correlative with Fab melting temperature. We aim to develop a model whose output is predictive of T_m_, while maintaining the speed with which we can currently generate predictions based on covariance violations. Based on previous literature, we hypothesize that embeddings generated from PLMs fine-tuned on paired antibody variable domains (e.g., IgBERT) contain information that is relevant to protein stability.^31,38,39^ To test this hypothesis, prior to any additional supervised training of a model we first average each of the 1024 individual output embedding vectors over the protein length, then took the mean of those 1024 averages and plot T_m_ as a function of a single embedding mean to identify a correlation (Fig. 2a). This is what we refer to as a zero-shot prediction of melting temperature. We observe a weak correlation, a Pearson correlation coefficient 0.31, which was lower than our covariance baseline approach (Fig. 2a, Table 2).

**Figure 2:**
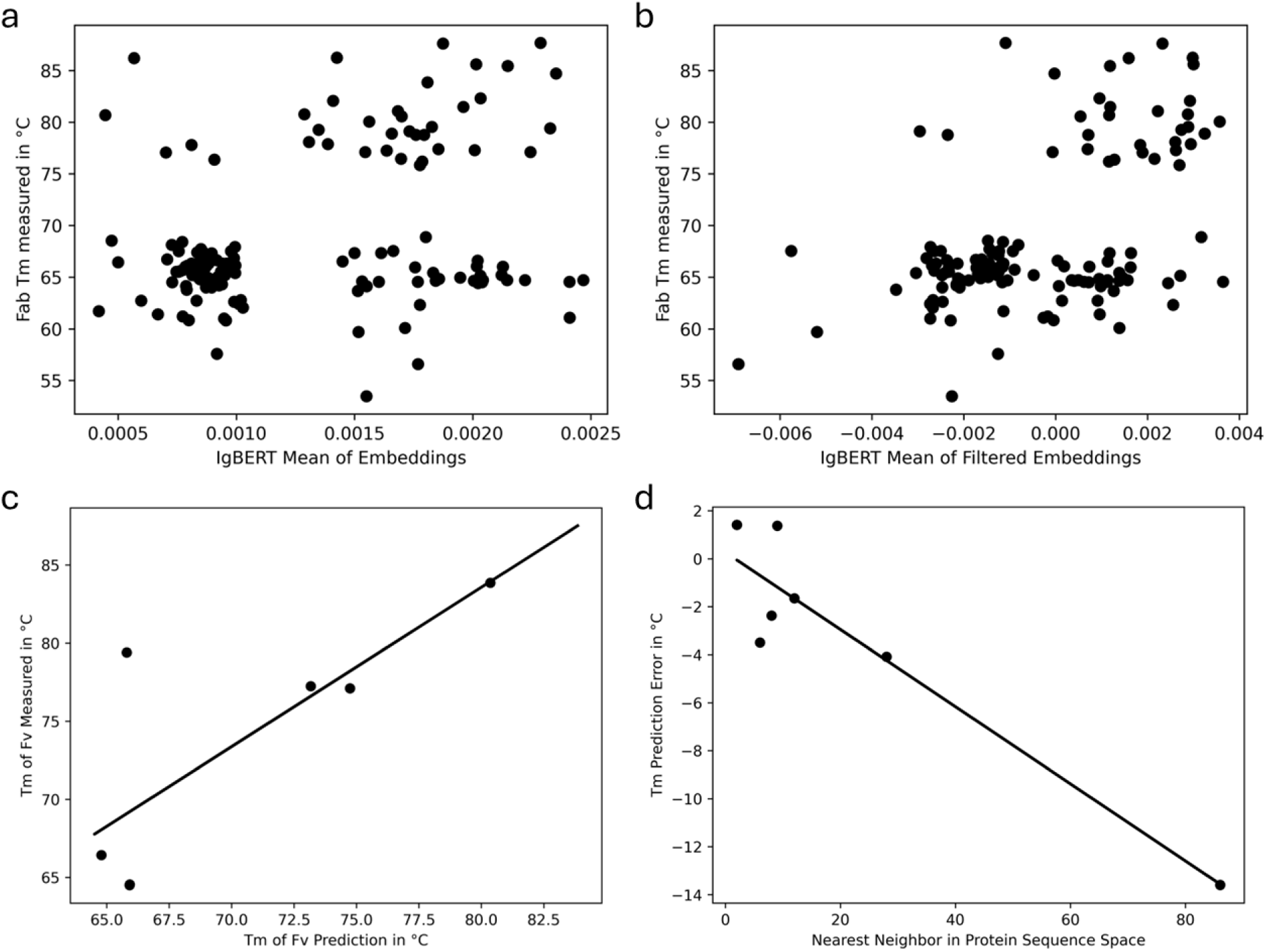
IgBERT is able to capture features that are relevant and predictive of melting temperature, (a) A scatter-plot of Fab T_m_ as a function of the mean of the global embeddings averaged over the length of the Fv sequence, (b) A scatter-plot of Fab T_m_ as a function of the mean of a subset of the embeddings averaged over the length of the Fv sequence filtered by a Pearson correlation coefficient of |0.5|. (c) Random Forest Regressor model trained to predict Tm of the Fab using IgBERT embeddings, (d) Error in T_m_ prediction test set as a function of amino acid differences from nearest neighbor in the training set.

**Table 2:** Performance of T_m_ predictions using the IgBERT embedding.

|  | slope | intercept | PCC |
| --- | --- | --- | --- |
| Zero Shot – All | 4191.31 | 63.34 | 0.31 |
| Zero Shot - Filtered | 1770.94 | 69.20 | 0.50 |
| RF Regressor | 1.02 | 1.97 | 0.77 |
| Regressor Error | -0.16 | 0.27 | -0.94 |

We hypothesized that there are some embeddings that do not correlate with the melting temperature. Furthermore, some proportion of the embeddings are likely correlate to each other providing redundant information. If this is true, then removing uncorrelated and redundant embeddings would improve our zero-shot T_m_ correlation. To address this, we filtered the embedding output from IgBERT first by correlation to T_m_ and second by correlation with remaining embeddings (SI Fig. 1). This reduced the IgBERT embeddings from 1024 down to 177. As we expected, the Pearson correlation coefficient improved from 0.31 to 0.50 (Fig. 2b, Table 2). This outperformed our baseline covariance analysis.

We infer that the 177 embeddings capture some relevant information with respect to the Fab T_m_, given the correlation we observed in our zero-shot analysis. We reasoned these could be used to train a supervised model to predict Fab T_m_. So, the T_m_ correlated embeddings from IgBERT were used to train a random forest regressor with the goal of predicting the Fab T_m_. The model was tested on a held-out internal dataset of 7 Fabs (Fig. 2c). A linear least-squares regression was performed on the measured Fab T_m_ as a function of predicted Fab T_m_. The Pearson correlation coefficient was 0.77 with a slope of 1.02 and an intercept of 1.97 (Table 2).

A known problem with ML models is the inability to generalize to unseen data. Is the reason most predictions are accurate because they are close in sequence space to a molecule in the training data? To answer this question, we also plot the difference between T_m_ prediction and truth as a function of nearest neighbor protein sequence (Fig. 2d). The closer the test sequence from the Jain set is to a training sequence in our internal dataset the better the prediction. This points toward our model having limited generalizability, but given the small test set it is unclear how limited our model is.

### Our ML Model outperforms our statistics-based model for the validation dataset

Like with our statistics-based model, our ML model is correlative with our internal dataset melting temperatures. However, does our ML model continue to be informative on the melting temperatures of Fabs in the publicly available dataset, unlike with our statistics-based model? The expectation is that our predictions will get worse, but still on average allow us to differentiate those Fabs that are likely to be more stable from Fabs that are likely to be less stable. Indeed, our predictions become less accurate and biased for the publicly available dataset (Fig. 3a, Table 3). The Pearson correlation coefficient is 0.28 between predicted and measured T_m_. We observe a clear reduction in the ability of our model to predict melting temperatures of Fab sequences. Nevertheless, our ML model clearly outperforms our statistics-based model and sequence-only approach.

**Figure 3:**
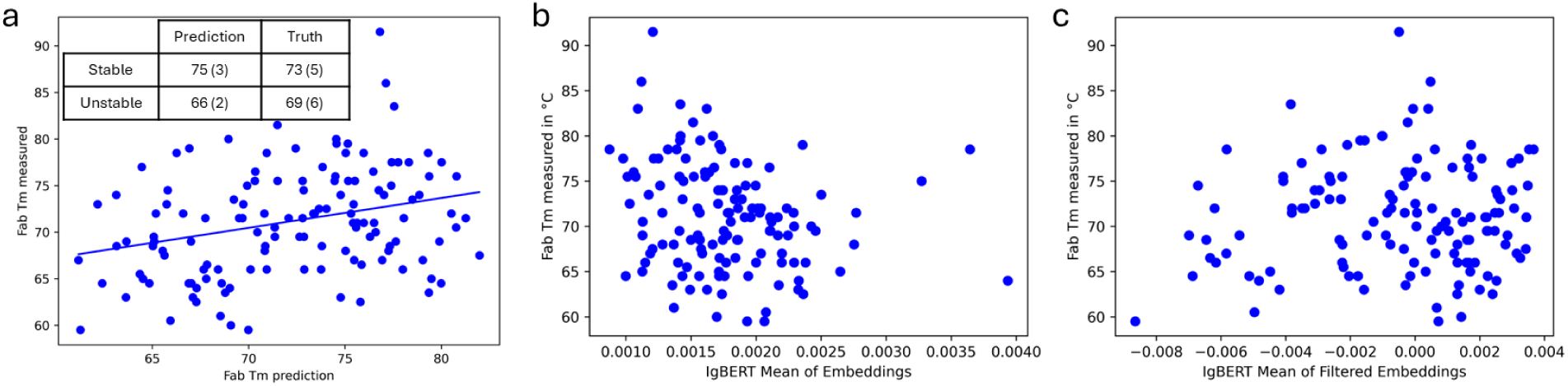
The trained model is more predictive of thermostability than our baseline model, (a). Random Forest Regressor model trained on our internal dataset used to predict T_m_ of the Fab for clinical stage therapeutics. Inset shows the mean and standard deviation for sequences predicted to be above and below 70 °C for the predicted and true T_m_. (b) A scatter plot showing the mean of all embeddings that were globally averaged over the input length of the Fv Sequence, (c) Is the same as (b), but the mean of the embeddings was filtered based on the training set correlations.

**Table 3:** Comparison of T_m_ prediction between our benchmark model and IgBERT embedding derived models.

|  | slope | intercept | PCC |
| --- | --- | --- | --- |
| RF Model | 0.32 | 48.00 | 0.28 |
| Zero Shot – All | -2738.96 | 75.97 | -0.23 |
| Zero Shot - Filtered | 138.36 | 71.22 | 0.07 |

The ultimate goal is to differentiate a set of Fabs based on thermostability, not necessarily predict the T_m_. If we consider Fabs with a predicted T_m_ above 70 °C as stable and those with a predicted T_m_ below 70 °C as unstable, then we can simplify the task. The null hypothesis is that our model is unable to differentiate stable from unstable Fabs. More specifically, the true mean of the unstable distribution is equal to the true mean of the stable distribution. The true mean of the Fabs predicted to be stable is 73 (±5) °C, which is greater than the true mean, 69 (±6) °C, of the Fabs predicted to be unstable (Fig. 3a (inset)). A p-value of 3.68*10^-4^ was calculated using Welch’s t-test to account for unequal variances, which indicates we can likely reject the null hypothesis that the two distributions are equal. Consequently, we can, on average, differentiate Fabs based on thermostability.

Our model retains some predictive power for the clinical stage therapeutic dataset but is reduced. One potential explanation for this is that the Jain set are different in some way from the internal dataset we have aggregated. We plot the Fab T_m_ as a function of the mean of all embeddings output from IgBERT (Fig. 3b). Unlike with our internal dataset, the majority of the clinical stage therapeutic mean of embeddings are >0.00125. In addition, they even range up to 0.004 where the internal dataset only goes up to 0.0025. When we plot the Fab T_m_ as a function of the mean of the 177 embeddings filtered based on our internal dataset correlation analysis, then we see that the range of embedding values for the Jain set is similar to the internal dataset (Fig. 3c). We infer that we have removed features that differentiated the Jain set from internal dataset but did not necessarily impact T_m_.

### Training-set and test-set embedding differences limit model generalizability

In an actual application of the model, the sequences we are given to predict thermostability will almost always be unique as compared to our training data. Therefore, a good feature of a test set is that it is different from the training data. To quantify this difference, we performed t-SNE on the filtered PLM embeddings of the training set and the Jain set (Fig. 4).

**Figure 4:**
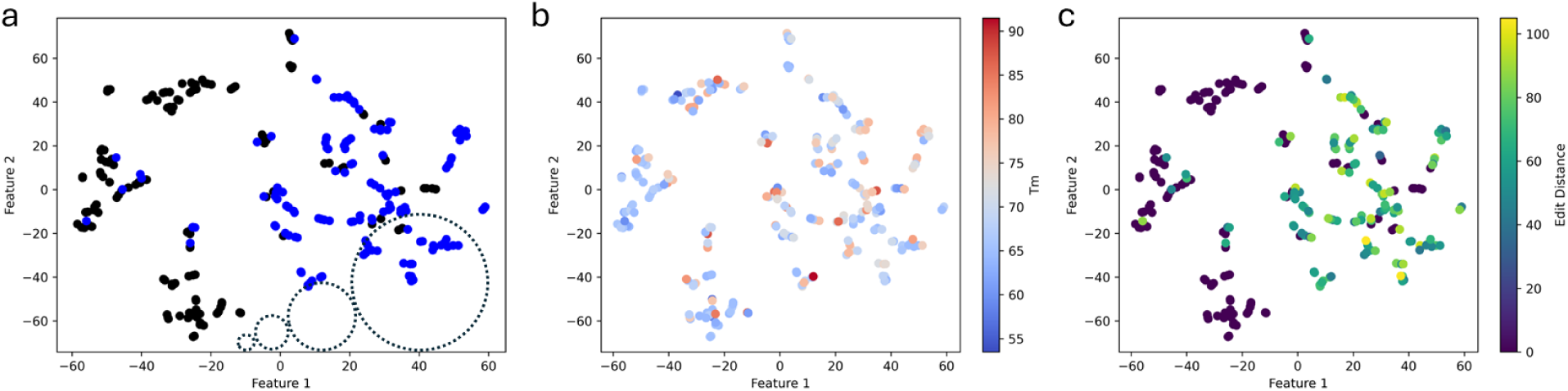
The distribution of embeddings for test set is different from the training set. (a). A t-SNE of the training (black) and test data (blue) together. The four dotted-circles represent a cartoon of the cutoff area if a training point Lay at the center. The radii are 2.5, 5,10, and 20 from left to right, (b) & (c) are the same as (a), but with each point colored as a function of the Tm or sequence differences from the training set nearest neighbor respectively.

From Fig. 4a we can see that the majority of the test set clusters separately from the training data. The distribution of embeddings output for the Jain set is distinct from the embeddings output for the training set.

The t-SNE features for training and Jain sequences cluster among themselves. Previous research has shown that t-SNE representations tend to cluster together based on distance in a phylogenetic tree, sequence identity, and other attributes.^25,28,33,40–42^ Do we see the same clustering for T_m_ or edit distance in our analysis? To test this, each molecule was colored according to its melting temperature (Fig. 4b) or shortest edit distance from a sequence in the training set (Fig. 4c). We do not see sequences with a high T_m_ (red dots) clustering among other sequences with a high T_m_. Rather, sequences with a high T_m_ are interspersed among sequences with low T_m_ (blue dots) illustrating the difficulty of T_m_ prediction. In addition, different sequences with different edit distances from an internal sequence cluster together.

The difference between the training set and Jain set embedding distribution does not appear to be a function of T_m_ or sequence identity.

Despite no obvious clustering between t-SNE features and T_m_ or sequence identity, we expect a correlation still exists. We hypothesize that if there is a correlation between PLM embedding distributions and T_m_, then model predictions will improve with decreasing t-SNE feature distance between a training and Jain sequence. To test this hypothesis, we created 4 arbitrary radial thresholds and looked at true T_m_ as a function of T_m_ prediction (Fig. 5). As we expected, the closer in embedding space the test set is to the training set the more accurate the model T_m_ predictions.

**Figure 5:**
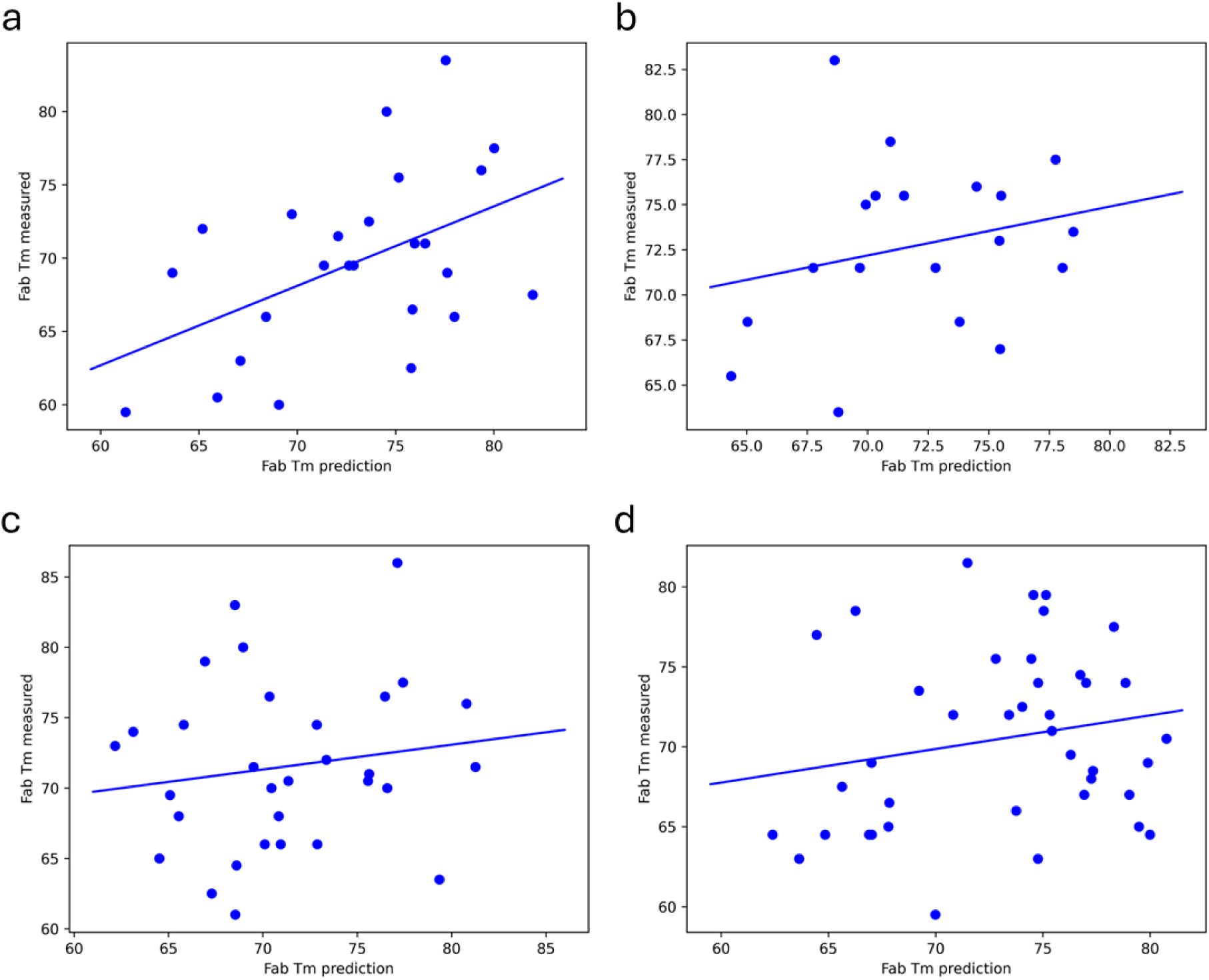
Restricting sequence T_m_ predictions to those similar to the training set, as defined by t-SNE, increases confidence in our predictions, (a). A scatter plot of the true Fab T_m_ as a function predicted Fab T_m_ from our model at a t-SNE distance range of [0, 2.5) to the nearest neighbor in the training set. (b) -(d) are the same as (a) but with ranges of [2.5, 5), [5, 10), and [10, 20) respectively.

Placing a restraint on sequences within the test set to be within a radius of 2.5 to the nearest training point increases the Pearson correlation coefficient value from 0.28 to 0.48, while still including almost 20% of the test set (Table 4). T_m_ predictions for molecules within the (2.5, 5] range have a Pearson correlation coefficient value of 0.24 and includes an additional 14% of the test set (Fig. 5b). Predictions beyond these ranges display very little correlation with ground truth (Figs. 5c & 5d). For each bin, the average edit distance of a test sequence from the closest training sequence (neighbor string distance) does not change monotonically (Table 4). The closer the distribution of a test sequence embedding is to a distribution of a training sequence embedding, and not necessarily sequence space, the closer our prediction is to ground truth.

**Table 4:**
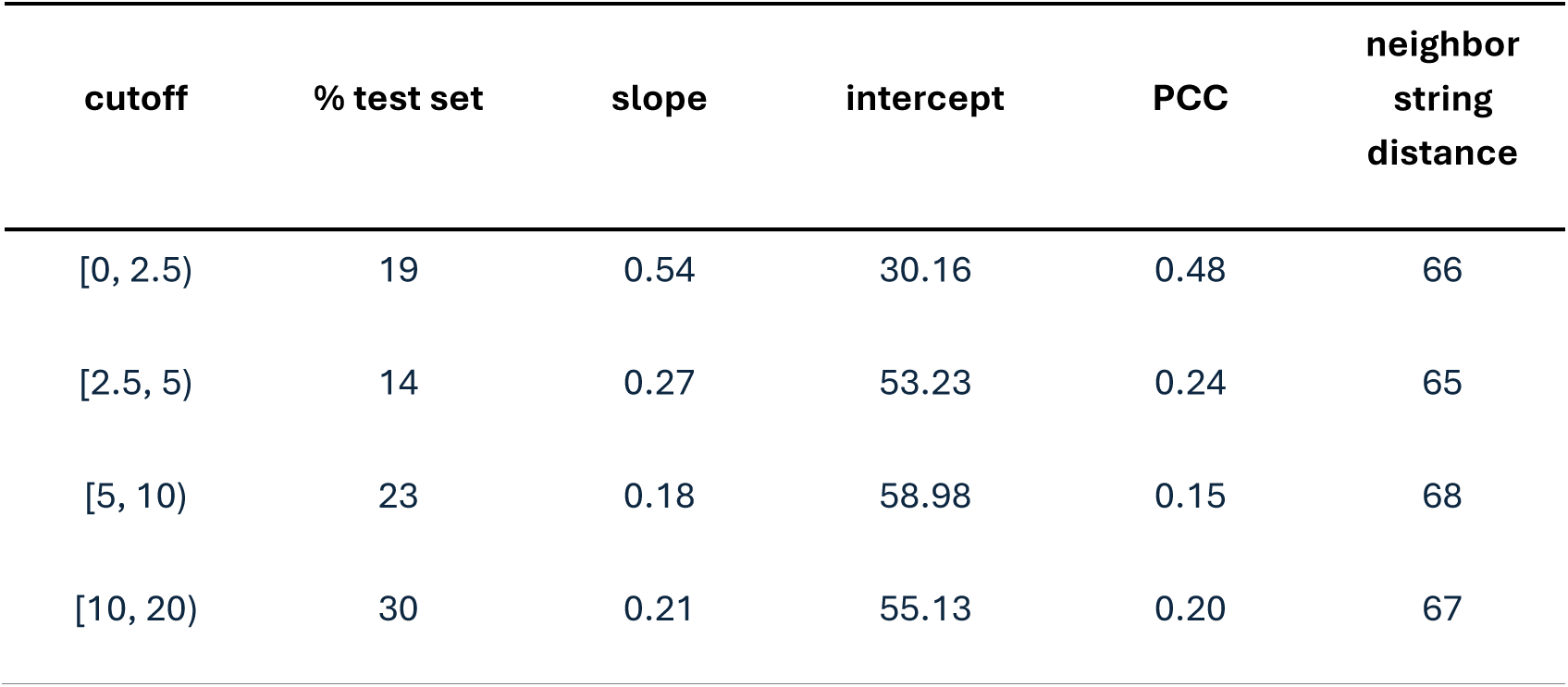
Comparison of model performance as a function of IgBERT embedding differences.

| cutoff | % test set | slope | intercept | PCC | neighbor<br>string<br>distance |
| --- | --- | --- | --- | --- | --- |
| [0, 2.5) | 19 | 0.54 | 30.16 | 0.48 | 66 |
| [2.5, 5) | 14 | 0.27 | 53.23 | 0.24 | 65 |
| [5, 10) | 23 | 0.18 | 58.98 | 0.15 | 68 |
| [10, 20) | 30 | 0.21 | 55.13 | 0.20 | 67 |

## Discussion

PLMs output a vector of embeddings that are expected to contain information relevant to the biochemical and biophysical properties of proteins. These embeddings are the result of learning contextual information at each site of a protein. Can this contextual information provide any insight into specific biophysical properties for a specific class of protein? We demonstrate that these models do in fact learn information relevant to predicting melting temperatures of the antigen binding fragment of an antibody. While the model cannot exactly predict T_m_, overall, the model can still distinguish thermally stable Fabs from thermally unstable Fabsquickly and cheaply for a publicly available test set. This is a scalable tool to de-risk lead antibody candidates based on conformational thermostability.

### Sequence-based statistical models are limited

A sequence only approach to predicting thermostability is desirable because it is scalable for lead candidate selection. Initially, we used covariance analysis to differentiate molecules based on thermostability. Covariance analysis, and similar methods, has been demonstrated to identify pairs of residues contacting each other in the tertiary structure of a protein from primary structure alone.^15,17–20,43^ Our expectation is that disrupting pairs of interacting residues is likely to disrupt the protein fold and reduce conformational stability. As a result, the T_m_ of the proteins with more covariance violations will be lower than proteins with fewer covariance violations.

Covariance analysis provides a simple and interpretable method to differentiate antibodies based on thermostability. For example, burying a hydrophilic or charged residue in the protein core when covarying sites are hydrophobic would be considered detrimental. We see that for our internal datasets, an increase in the number of covariance violations present in an antibody sequence correlates with lower thermostability (Fig. 1).

However, when we extend this analysis to the Jain set a weak positive correlation is observed (Fig. 1c). This observation is at odds with our interpretation of the model and highlights the model limitations that there is no scenario in which introducing a mutation at a covarying site will enhance stability. In reality, there is likely a subset of sites where changing the chemical nature of a residue pair can be beneficial. Therefore, we needed a model that could consider mutations as beneficial to stability rather than only detrimental.

### Our ML model Fab T_m_ predictions outperform a baseline statistical-model

The correlation between our covariance analysis and T_m_ fails to extend to our test set and there is no correlation between test Fabs closest in sequence space to the training set (Fig. 1). Therefore, we turned to a supervised machine learning approach. However, this approach introduced two problems. First, we need a way to represent a sequence and its inherent information as a differentiable vector. Second, we need a lot of diverse data labelled with melting temperatures. We addressed the first problem by turning to a fine-tuned protein language model, IgBERT and the second problem by collating data from historical programs.

Several linear regression models were trained on the task to predict the Fab melting temperature given a set of filtered IgBERT embeddings derived from the paired Fv sequence. A Random Forest model was selected according to PCC, and the slope and intercept from a linear least squares regression between predicted T_m_ and actual T_m_. Our best model yielded a PCC of 0.77, a slope of 1.02, and an intercept of 1.97. To determine how well our model would generalize, we predicted the Fab T_m_ for over 130 molecules in the Jain set (Fig. 3a). A regression analysis of the predicted and true T_m_s resulted in a PCC of 0.28, a slope of 0.32, and an intercept of 48 (Table 3). Despite this decrease in model predictiveness, it still outperforms the covariance analysis.

### IgBERT embeddings contain information relevant to Fab T_m_ prediction

A PLM is a transformer-based architecture that is expected to learn correlations among amino acids in a protein sequence through a masked language objective. The output of these models are embeddings ─ which are set of vectors ─ whose value is expected to be dependent on the model learning relevant contextual relationships of protein tertiary and secondary structure in its task to recover protein primary structure. Given a trained PLM, the embeddings are cheap and fast to generate, making them attractive features to train supervised ML models on specific tasks.

IgBERT is an antibody specific PLM that is ProtBERT fine-tuned on unpaired and paired OAS sequences. It is one of the top-performing antibody-specific PLMs with respect to sequence recovery. Given that, we chose to use IgBERT to generate paired Fv embeddings as features to train a supervised model on a thermostability prediction task. However, prior to training a model we first chose to assess whether there was any correlation between IgBERT Fv embedding and melting temperature (Fig. 2a). We observed that if the mean embedding was < 0.00125 the protein sequences were likely to have a lower T_m_ than if the mean embeddings were > 0.00125. We infer that the embedding set for each Fv contain some relevant information with respect to protein thermostability and it is worth training a downstream model on a thermostability prediction task.

There are 1024 embeddings globally averaged over the length of the protein sequence derived from IgBERT. While each embedding is a function of the entire length of the input protein, we reasoned a subset of those embeddings are more predictive of T_m_ than others. For example, we expect that some embeddings are more heavily influenced by the highly variable CDRs. This would likely introduce noise that a downstream model would need to learn to filter out, thus increasing the difficulty of thermostability prediction.

Therefore, before training a model we identified a subset of embeddings that correlated to melting temperature and did not contain redundant information (SI Fig 1). This reduced the number of embeddings from 1024 to 177. We then plot the T_m_ as a function of the average of the 177 embeddings (Fig. 2a & 2b). The Pearson correlation coefficient increases 0.19 for the training set. This improvement demonstrates we are removing factors that could confound a downstream model in predicting T_m_.

### Filtering IgBERT embeddings based on Fab T_m_ improves average similarity between the training and Jain set

By filtering embeddings based on their correlation to the melting temperatures of our internal training dataset we run the risk of reducing predictive power to an external test set. We see that the distribution of the mean for training dataset embeddings is different from the distribution of the mean for test set embeddings (Figs. 2a & 3b). For our test set, the mean unfiltered embedding values for all Fv sequences is greater than 0.00125, whereas a substantial portion unfiltered embedding mean values of unstable Fv sequences are below that threshold. However, after the embeddings are filtered by PCC with the melting temperature of our internal dataset, we observe the distribution of mean of embeddings for the test set better reflect the training set (Figs. 2b & 3c).

Why do we observe the mean embedding value of the test and training dataset become more similar post-filtering? As we hypothesized earlier, each embedding is a function of the entire length of the input protein but some of those embeddings are likely to be more informative toward other biophysical and biochemical characteristics of the Fv sequence. Fundamentally, other biophysical and biochemical properties (e.g., antigen binding specificity) may have exaggerated the difference between the training and Jain sets.

### Embedding similarity, not sequence identity, drives accurate model predictions

If a model is to be useful, then we must understand the limitations of that model. Otherwise, we run the risk of misuse. With that in mind, how do we know when we are appropriately applying our model to a molecule for prediction? Our expectation is that if we see a set of new molecules that is similar to the training data then our model would be more likely to differentiate that new set on the basis of thermostability. To accomplish this, we first needed a method to determine similarity among molecule sets.

One option for quantitatively determining similarity among molecule sets is t-SNE. In the past, t-SNE has been shown to reduce embedding distributions to cluster based on associated attributes. We reasoned that test sequences with similar embedding distributions to training sequences are likely to have more accurate predictions. The t-SNE analysis was used to reduce our input features from 177 to 2 dimensions (Fig. 4). After dimensionality reduction to two features, we clearly see that embedding distributions for a majority of the Fv sequences from the Jain set are distinct from the training set with some Jain set molecules being closer in Euclidean space to training set molecules than others.

The expectation is that predictions for sequences in the test set near our training data will be more accurate than those further away. This is in fact what we see; the closer the t-SNE embedding value of the test set is to the training set the more accurate the model prediction (Fig. 5). The Pearson correlation coefficient of predicted melting temperatures to actual melting temperatures is 0.48 when test points are within a radius of [0, 2.5) (Table 4). There is a drop in the Pearson correlation coefficients to a range of 0.15 to 0.24 for data further away. Crucially, edit-distance is essentially uncorrelated with model performance.

Why does edit-distance not correlate with accuracy of our model predictions? One reason is that edit-distance is potentially an overly simplistic metric. Sequences can have the same edit-distance, but radically different outcomes. For example, introducing a different hydrophobic variant buried in the protein will be less destabilizing than introducing a charged residue. Another reason is that our model is trained directly from embeddings of a PLM rather than the actual Fv sequence itself. While sequence determines the embedding vector it is not a simple linear transformation. Therefore, we reason that if model performance is to be improved in subsequent iterations it is likely important to sample the diversity of embedding space used to train the model, in addition to sequence diversity.

### Our model is able to de-risk the manufacturing process for a set of lead candidates

The original goal was not necessarily to accurately predict the Fab T_m_ but to be able to differentiate Fabs based on thermostability to de-risk the manufacturing process. Practically, these tools can be implemented in different ways to achieve an ultimate goal of eliminating lead candidates with properties that would stop or delay their manufacture. One approach, analogous to the method by Raybould et al., is to down select a panel of lead candidates by setting a threshold in which an attribute or property is considered acceptable.^10^

The down-selection approach could be applied to T_m_. The acceptable threshold may be anything above 70 °C because this is the approximate melting point of the Fc domain. Increasing stability above this point may result in diminishing returns on investment. We find that the average melting temperatures of Fab T_m_ predicted to be stable (i.e., above 70 °C) by our model is 73 (±5) °C. Assuming a normal distribution, roughly 70% of the proteins predicted to be stable are in fact stable. Whereas random sampling or covariance would essentially be a coin flip.

## Conclusion

Differentiating many antibody lead candidates based on thermostability quickly is a difficult task. We train and task a model to differentiate Fv domains based on their T_m_ with limited success. We demonstrate that preprocessing the embeddings results in a subset of embeddings that better correlate to thermostability. Furthermore, we demonstrate what impacts model accuracy is the difference between training embedding distribution and Jain embedding distribution. Nevertheless, on average our model is able to segregate Fv domains based on thermostability to de-risk the manufacturing process and there is a subset for which the predictions display a higher correlation to ground truth. We present a scalable alternative to differentiating antibodies based on thermostability based solely on Fv domain sequence.

## Acknowledgements

I would like to thank all those at Just-Evotec Biologics who acquired the data and had the foresight to record it in the same excel format.

