## Supplementary material for "Sequence-Only Prediction of Antibody Fab Thermostability Using Protein Language Model Embeddings": Antibody_Thermostability_Predictions_SI_v4.1.pdf

### Supplemental Figures

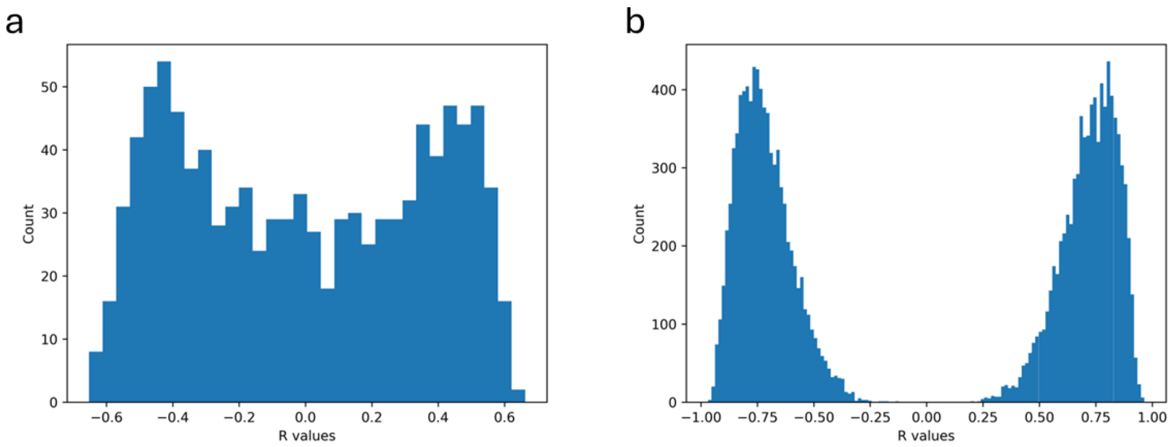


SI Figure 1: A fraction of embeddings uniquely correlate to thermostability for our internal dataset. (a). A histogram of correlation between sequence embedding and Fv Tm. (b) A histogram of correlation between embeddings filtered from (a) based on a threshold that R value be above 0.5 or less than -0.5.

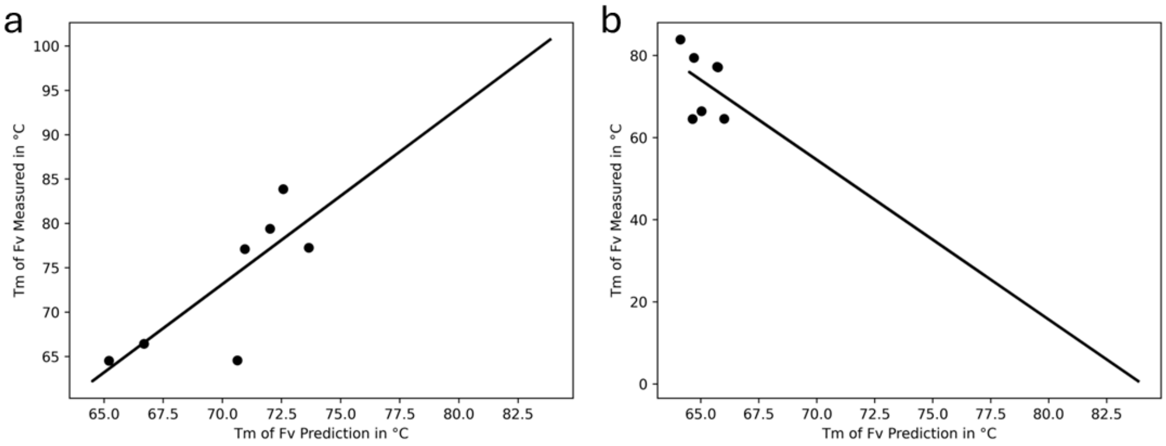


SI Figure 2: Training additional models offers little to no improvement on Fab Tm prediction. (a). A scatter plot of actual Fab Tm as a function of predicted Fab Tm using an SGD regressor. (b) The same as (a), but using a SVM regressor.

| | slope | intercept | PCC |
|---|---|---|---|
| RF | 1.02 | 1.97 | 0.77 |
| SGD | 1.99 | -66.16 | 0.78 |
| SVM | -3.89 | 326.89 | -0.34 |

SI Table 1: Comparison of model performance for internal dataset.

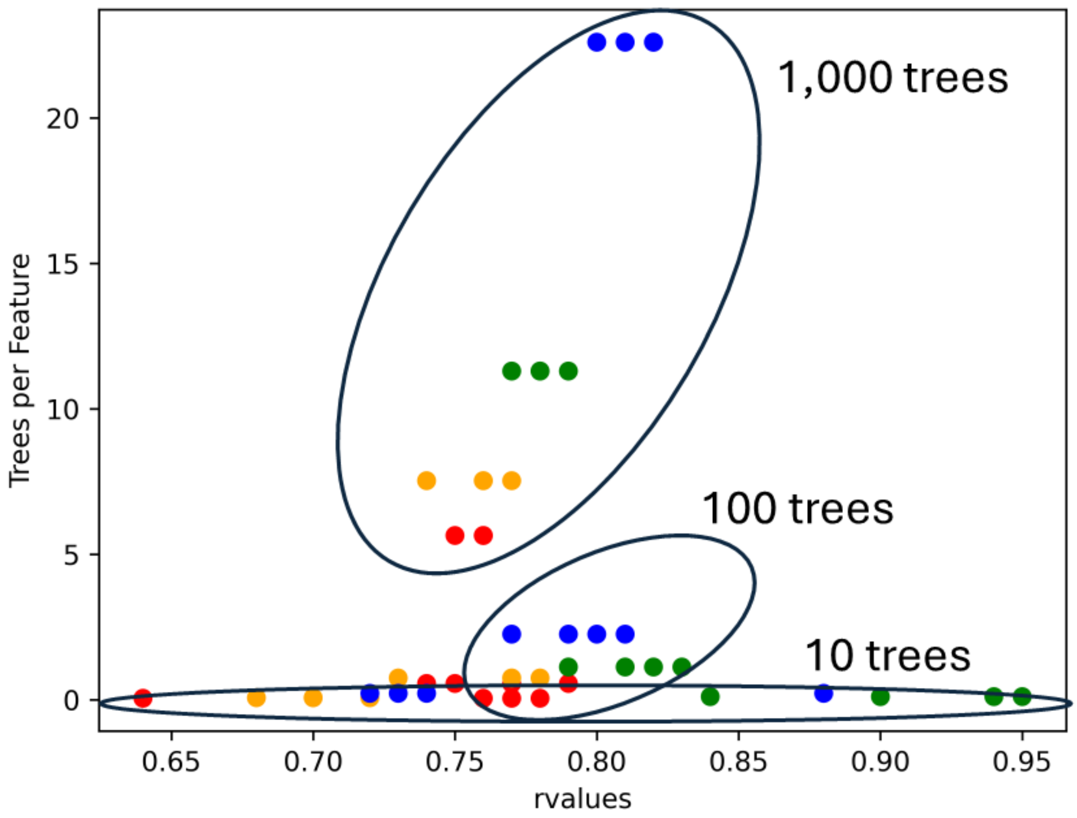


SI Figure 3: Hyperparameter search of Random Forest regressor yielded models with an increase in Pearson correlation coefficients. This is a scatter plot of showing Trees per Feature as a function of Pearson correlation coefficient. The markers on the graph are colored by the fraction of embeddings used to train a model. Blue, green, orange, and red represent 0.25, 0.5, 0.75, and 1.0 respectively.

| Data Usage | Trees | Scoring Fxn | Sum(Pred – True) | PCC | <Std> |
|---|---|---|---|---|---|
| 100% | 100 | *Squared Err* | *-22* | *0.77* | *5.3* |
| | | Abs Err | -22 | 0.74 | 5.1 |
| | | MSE | -21 | 0.75 | 5.3 |
| | | Poisson | -22 | 0.79 | 5.3 |
| | 10 | Squared Err | -21 | 0.77 | 5.0 |
| | | Abs Err | -28 | 0.64 | **<u>3.8</u>** |
| | | MSE | -21 | 0.76 | 4.9 |
| | | Poisson | -23 | 0.78 | 4.9 |
| 50% | 100 | Squared Err | -23 | 0.81 | 5.4 |
| | | Abs Err | -22 | 0.79 | 5.2 |
| | | MSE | -24 | 0.83 | 5.4 |
| | | Poisson | -24 | 0.82 | 5.6 |
| | 10 | Squared Err | -24 | 0.94 | 5.8 |
| | | Abs Err | **<u>-12</u>** | 0.84 | 5.6 |
| | | MSE | -25 | **<u>0.95</u>** | 5.5 |
| | | Poisson | -34 | 0.90 | 6.0 |

SI Table 2: Comparison of different Random Forest models with different hyperparameters. Bolded and underlined are models with the highest values for that metric.

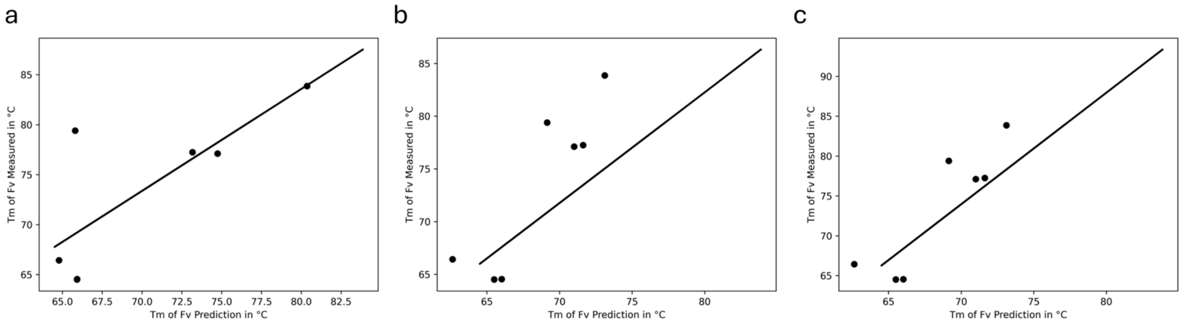


SI Figure 4: Best results from hyperparameter search demonstrate increases in Pearson correlation coefficient coincide with an increase in model prediction bias. (a). A scatter plot of actual Fab Tm as a function of predicted Fab Tm for our initial Random Forest Regressor model. (b) and (c) are the same as (a) but using two different metrics for success. (b) The best model is selected based on the sum of predictions being closest to 0. (c) The best model is selected based on the Pearson correlation coefficient.

| | **slope** | **intercept** | **PCC** |
|---|---|---|---|
| RF-0 | 1.02 | 1.97 | 0.77 |
| RF-Sum Err | 1.05 | -1.77 | 0.84 |
| RF-R | 1.40 | -24.05 | 0.95 |

SI Table 3: Comparison of the top 3 Random Forest models

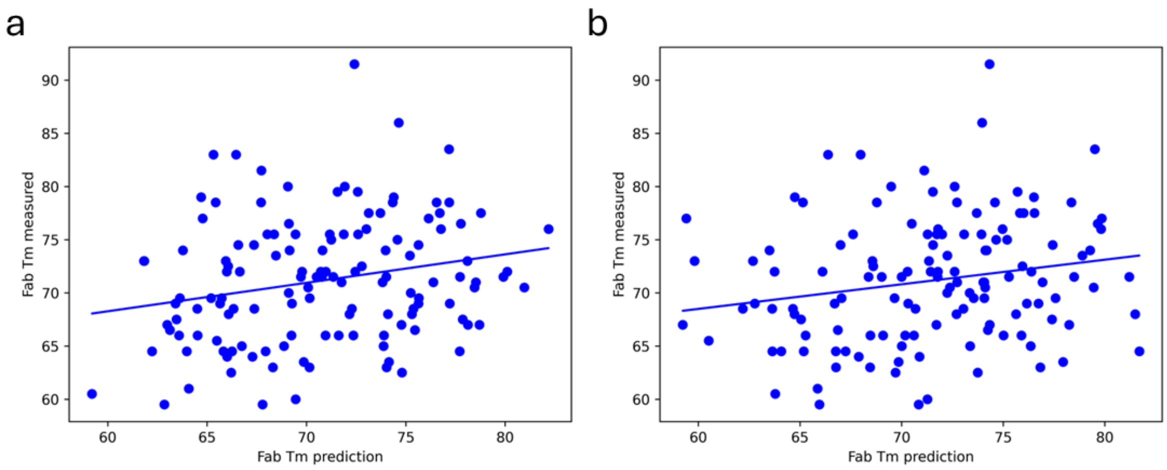


SI Figure 5: Better Random Forest models, as judged by sum or errors or Pearson correlation coefficient, do not in fact result in more predictive models for unseen data (a). A scatter plot of actual Fab Tm as a function of predicted Fab Tm for our best model as judged by Pearson correlation coefficient. (b) The same as (a) but using the best model based on the lowest sum prediction errors.

| | **slope** | **intercept** | **PCC** |
|---|---|---|---|
| RF-0 | 0.32 | 48.00 | 0.28 |
| RF-sum Err | 0.23 | 54.62 | 0.20 |
| RF - R | 0.27 | 52.2 | 0.23 |

SI Table 4: Comparison of the best Random Forest models performance for the publicly available dataset.
